# Yellow fever vaccine propagation in primary human hepatocytes triggers antiviral and cytolytic responses

**DOI:** 10.64898/2026.01.21.700765

**Authors:** Robin D.V. Kleinert, Maike Herrmann, André Gömer, Leyla Sirkinti, Csaba Miskey, Yannick Brüggemann, Nadi Dixit, Theresa S. Bechtel, Maximilian Beikirch, Nicola Frericks, Bingqian Qu, Bevan Sawatsky, Olga Tichá, Julia Hehner, Yvonne F. Grande, Eva Herker, Joe Grove, Eike Steinmann, Daniel Todt, Richard J.P. Brown

**Affiliations:** Division of Veterinary Medicine, Paul-Ehrlich-Institut, 63225 Langen, Germany; Host-Pathogen-Interactions, Paul-Ehrlich-Institut, 63225 Langen, Germany; Department of Molecular and Medical Virology, Ruhr University, 44801 Bochum, Germany; Department of Translational and Computational Infection Research (TRACiR), Ruhr University, 44801 Bochum, Germany; Genomics Core Facility, Paul-Ehrlich-Institut, 63225 Langen, Germany; Division of Infectology, Paul-Ehrlich-Institut, 63225 Langen, Germany; Institute of Virology University of Marburg, 35037 Marburg, Germany; MRC–University of Glasgow Centre for Virus Research, Glasgow, UK

## Abstract

Yellow fever virus (YFV) infection can cause severe-to-fatal liver damage in humans, while immunization with the attenuated 17D vaccine strain has an excellent safety record, priming protective host immunity in the absence of pathology. To investigate virus-host correlates associated with these differential clinical outcomes, we combined YFV genome-level evolutionary analyses with investigations of vaccine and virulent strain hepatotropism. Evolutionary analyses confirmed purifying selection is the dominant force driving global YFV genome divergence, with functional constraints associated with the YFV transmission cycle selecting against vaccine attenuating mutations in virulent strains. In immune deficient hepatoma cells, 17D exhibited enhanced early propagation, spreading and apoptosis induction when compared to virulent strains. *Ex vivo* infections performed in primary human hepatocytes (PHH) from multiple donors confirmed robust propagation of both 17D and virulent YFV strains. RNA-sequencing revealed consistent and shared induction of *IFNB* and *IFNL1-4*, modulating overlapping gene programs associated with antiviral responses, immunity, chemotaxis and inflammation, cell-death, metabolic reprogramming and protein translation. Subtle differences in virion production kinetics and the magnitude and tempo of PHH transcriptional responses were observed. Antiviral responses to 17D were activated earlier while responses to virulent YFV were delayed but enhanced, mirroring virus propagation kinetics. More broadly, cellular responses to YFV infection are likely dominated by paracrine IFN signalling, with enriched *LRP1* expression coupled with impaired IFNα production in PHH contributing to YFV’s robust hepatotropism. In summary, we demonstrate comparable propagation kinetics and PHH transcriptional responses to vaccine and virulent YFV strains, highlighting impaired hepatotropism is not a correlate of vaccine attenuation. These data imply that unknown barriers restrict liver access and associated organ pathology upon vaccination with 17D.

## Introduction

Yellow fever virus (YFV) is a member of the *Orthoflavivirus* genus and the prototypic member of the evolutionarily diverse and recently expanded family *Flaviviridae*, which includes a range of clinically relevant human and animal pathogens [1–3]. YFV is endemic in parts of tropical Africa and South America. The virus circulates in jungle primate reservoirs and is transmitted to humans primarily through the bite of infected *Aedes*, *Hamogogus* and *Sabethes* mosquitoes [4–6], with historical and contemporary outbreaks resulting in many fatalities [7–10]. Clinical disease manifests after an initial incubation period of 3-6 days, with symptoms including fever, headache, lower back pain, myalgia and bradycardia, which usually resolve after 3-4 days. In around 15-20% of cases, symptoms reappear and the disease progresses to a toxic phase characterized by bleeding from mucosal membranes, haemorrhagic shock and multi-organ failure. In these severe cases, mortality rates range from 20-50% [11,12]. While surveillance occurs in endemic regions, calculating the global burden of YFV is challenging. Mathematical modelling estimates 51,000 deaths and 109,000 severe cases attributable to YFV infections in both Africa and South America in 2018 [13]. While some YFV-targeting therapeutics have shown promise in experimental models, clinical management of YFV infected patients is often limited to treatment of symptoms [12]. However, a highly effective live-attenuated vaccine against YFV was developed in the 1930s and currently represents the most effective tool to limit YFV outbreaks [8,9,14].

The YFV vaccine was derived from a virulent patient isolate, the Asibi strain, and was attenuated by ∼200 serial passages, initially in mouse brains followed by minced chicken embryos with the brain and spinal columns removed [15]. This process resulted in the 17D strain, which exhibits a loss of virulence, associated visceral disease and mosquito competence, while replicative capacity and the ability to induce protective immunity is retained [16,17]. The YFV vaccine has an excellent safety profile and has been administered to over 850 million people globally [18], providing lifelong protection in most vaccinees [19,20]. The ∼11,000 bp YFV genome contains a single open reading frame (ORF) encoding a ∼3400 amino acid polyprotein, which is post-translationally processed by viral and host proteases into 10 mature proteins [21]. Compared to the parental Asibi strain, the 17D genome contains over 80 substitutions including 30 non-synonymous mutations [22,23]. While synonymous substitutions were shown to be dispensable for attenuation [24], non-synonymous mutations in the structural envelope (E) protein were demonstrated to enhance viral uptake and spread [24,25] while mutations in non-structural (NS) proteins NS2A and NS2B enhanced host antiviral responses and impaired virulence [24,26]. However, mechanistically, exactly how systemic attenuation in human vaccinees is achieved remains unclear.

YFV is unique amongst the *Orthoflavivirus* genus, with infections causing severe-to-fatal liver damage [11,27,28]. While many studies have investigated differences between the vaccine and the parental Asibi strain in tissue-culture and animal models, the virus-host correlates that differentiate attenuated 17D vaccine immunizations from liver-pathology causing virulent YFV infections are not well defined. With the aim of providing insights into vaccine strain attenuation, we first leveraged database deposited sequences and performed evolutionary analyses to evaluate selection pressures operating on virulent YFV genomes. We next explored YFV’s hepatotropism, with the aim of highlighting potential differences or similarities between vaccine and virulent strains. Comparative investigations were performed using authentic viral isolates: the 17D-204 vaccine strain (Stamaril^®^) and two genetically divergent virulent YFV strains. Cell-entry route, virus propagation kinetics and apoptosis induction rates were compared in human hepatoma cells that possess impaired innate immunity. Comparison of YFV strain-specific propagation kinetics and concomitant transcriptional responses to infection were also evaluated in explanted primary human hepatocytes (PHH), which are considered the gold standard cell model to study hepatic properties *in vitro* [29,30] and possess intact innate immunity, reacting strongly to viral infections [31–33]. Finally, liver entry-receptor expression, susceptibility to IFNα inhibition and the role of autocrine or paracrine IFN signalling in limiting YFV infection and spread were also investigated.

## Results

### Evolutionary analysis of globally sampled YFV genomes

For evolutionary analyses of virulent YFV genomes, inclusion criteria comprised a complete ORF, with subsequent removal of identical or vaccine strain genomes, resulting in a final panel of 272 sequences. First, pairwise comparisons of individual genomes and translated polyproteins were calculated and visualized as distance matrices (Fig. 1a). East African lineage (EAL) and Angola lineage (AL) viruses were distinct at up to 25% of nucleotide sites and up to 8% of encoded amino acid residues, from West African lineage (WAL) and South American lineage (SAL) viruses. Consistent with these analyses, phylogenetic reconstruction confirmed clustering into six distinct lineages, with significant bootstrap support (Fig. 1b). Consistent with an ancient separation ∼1000 years before present [34–36], a deep divergence was visible between the EAL/AL viruses and the WAL/SAL viruses. The distinction between WAL and SAL viruses, which represents ∼500 years of separation, was also clearly visible and strongly supported. Furthermore, these analyses also give an overview of recent YFV exports from endemic regions, highlighting virus transfer to Europe and Asia associated with globalized travel (Fig. 1b).

**Fig. 1.**
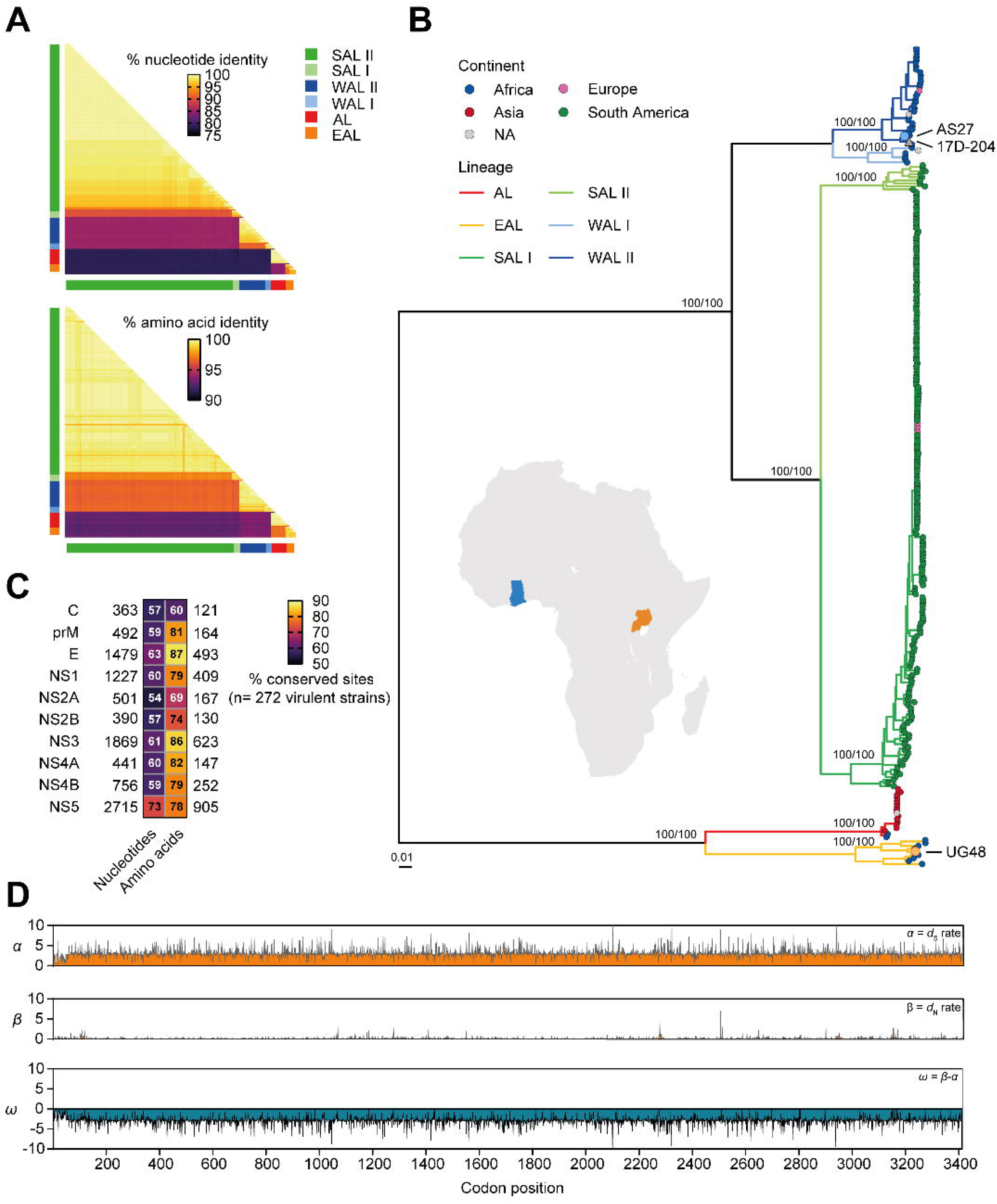
Purifying selection drives global YFV genome diversification. (*A*) Genome and encoded ORF comparisons of virulent YFV strains highlights six distinct YFV lineages. Pairwise distance matrix plots are derived from database deposited non-identical genomes from virulent isolates (n = 270), plus the genomes of virulent laboratory strains used in this study for downstream experiments and determined by NGS (n=2). Nucleotide (top panel) and amino acid (bottom panel) matrices represent pairwise comparisons of individual YFV genomes or encoded ORFs and are presented as % identity according to the scale bar. SAL: South American Lineage; WAL: West African Lineage; AL Angola Lineage; EAL: East African Lineage. (*B*) Geographical provenance and phylogenetic relatedness of globally sampled YFV strains. Phylogenetic analysis was performed with full-length ORF encoding sequences (10,236 – 10,239 bases) from virulent strains (n=272) plus the 17D-204 vaccine strain. Clades representing the six YFV lineages are colour-coded, with the geographical provenance of individual isolates highlighted at terminal nodes. Numbers assigned to internal tree nodes represent bootstrap support values derived from 1000 replications. Branch lengths are proportional to nucleotide substitutions per site, relative to the scale bar. Inset map of Africa highlights the sampling locations for virulent laboratory strains AS27 (Ghana: blue) and UG48 (Uganda: orange). (*C*) Nucleotide and amino acid conservation in individual YFV ORF encoded proteins. Heatmap displays conserved nucleotides (left) and amino acids (right) in individual YFV ORF encoded proteins from virulent strains (n=272). For each encoded protein, conserved sites are normalized (%) to total nucleotide or amino acid length, positioned to the left and right, respectively. (*D*) Purifying selection is the dominant force operating on the YFV ORF in globally sampled isolates (n=272). Top panel. Rate of synonymous substitutions at individual codon sites (*d*S = *α*) in the YFV ORF. Middle panel. Rate of non-synonymous substitutions at individual codon sites (*d*N = *β*) in the YFV ORF. Bottom panel. Normalized *d*N-*d*S rates at individual codon sites (*ω* = *β*-*α*) in the YFV ORF. Positive integers indicate diversifying (positive) selection, 0-values indicate neutral evolution, and negative integers indicate purifying (negative) selection.

To visualize potential differences in variability between protein-coding regions, levels of nucleotide and amino acid conservation in individual ORF-encoded proteins were calculated (Fig. 1c). Levels of nucleotide conservation were generally comparable across all genomic regions, although higher conservation was observed in NS5. Differences in levels of conservation were more apparent between individual encoded proteins. Of note, the highest level of amino acid conservation amongst all YFV ORF encoded proteins was observed in the envelope (E) protein, which mediates YFV entry into cells. These analyses highlight nucleotide variability in globally sampled genomes exceeds encoded amino acid variability (Fig. 1a and Fig. 1c), indicating functional constraints limit YFV protein evolution. Selection pressures which drive viral genome diversification and restrict amino acid variability likely differs at individual codons. To quantify these selection pressures, fine-scale codon-by-codon selection analysis scanning the entire YFV ORF was performed using FUBAR [37] (Fig. 1d). Synonymous substitution rates (*d*S = *α*: upper panel) far exceeded non-synonymous substitutions rates (*d*N = *β*: middle panel) across all codon sites. In line with this, no codon sites undergoing positive/diversifying selection were detected (*d*N - *d*S = *ω* >0: bottom panel). However, significant evidence for negative/purifying selection (*ω* <0) was detected at 3194 sites (posterior probability > 0.9), representing 94% of total YFV ORF codons. Together, these analyses highlight purifying selection as the dominant driver shaping global YFV genome diversification.

### Vaccine-specific mutations are selectively deleterious in virulent YFV

Comparison of the 17D-204 strain ORF with polyproteins from 272 virulent strains identified a constellation of 30 non-synonymous mutations correlated with vaccine attenuation (Fig. 2a). Focusing on sites at which vaccine-specific mutations occur, selection analysis of virulent strains confirmed a significant excess of synonymous over non-synonymous substitutions at these 28/30 codon triplets (*ω*<0, posterior probability >0.9) (Fig. 2a, upper panel). We next investigated whether vaccine-specific mutations circulate in globally sampled virulent YFV isolates. Frequencies of encoded amino acid residues at these 30 sites present in either attenuated 17D-204 or parental AS27 were calculated for our panel of virulent strains (Fig. 2a, lower panel). These data highlight strong amino acid conservation in virulent strains at these residues, with 17D-204-specific mutations either rarely or never observed at 29/30 sites. In summary, 28/30 codons at which vaccine-specific mutations occur exhibit significant signatures of strong purifying selection (an excess of synonymous over non-synonymous substitutions) and high levels of amino acid conservation in globally sampled virulent strains.

**Fig. 2.**
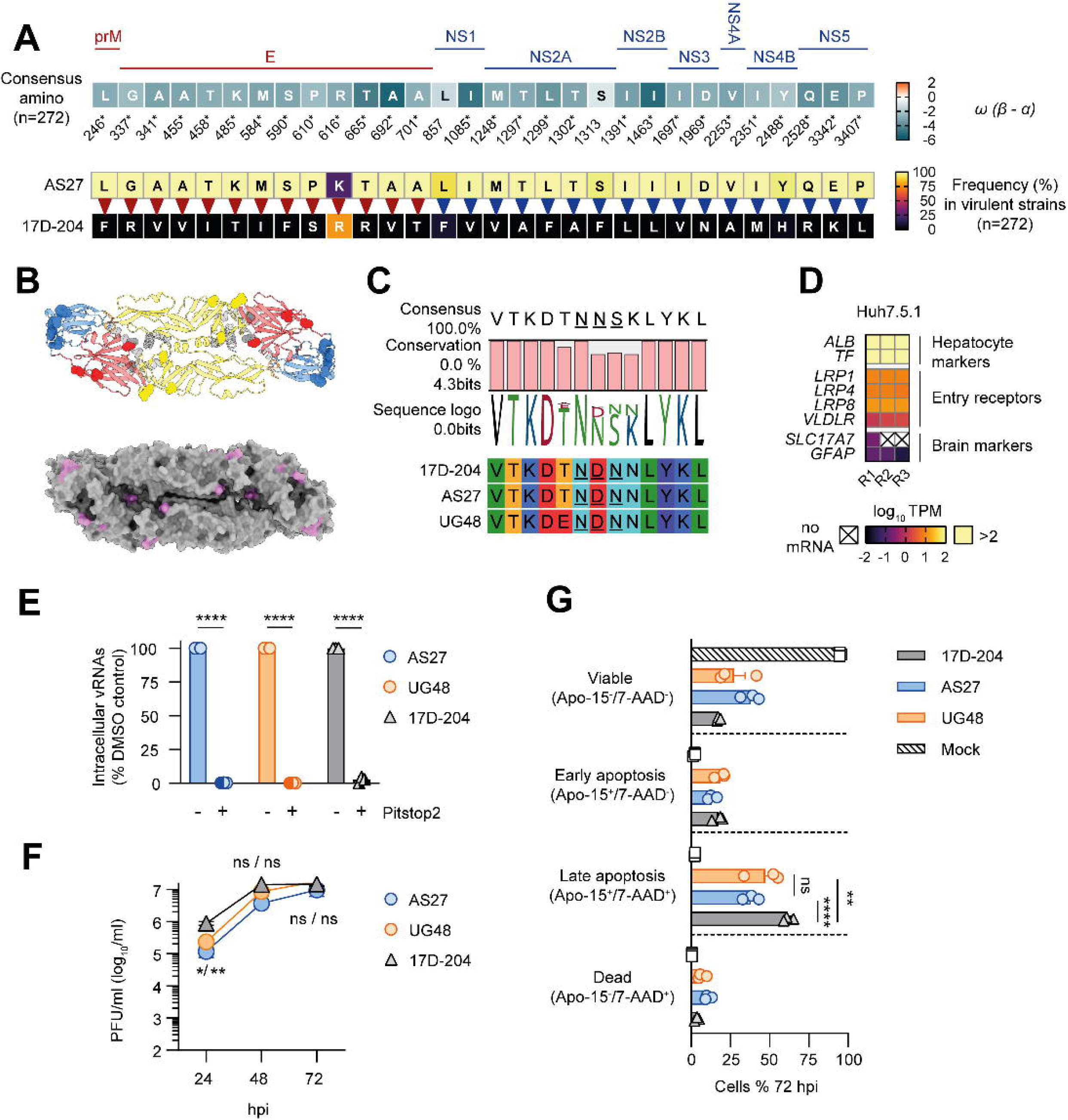
Vaccine-specific mutations are selected against in virulent YFV and confer enhanced propagation and apoptosis in immune deficient hepatoma cells. (*A*) Location, site-specific selection pressure and prevalence of 17D-204 specific mutations in virulent YFV strains. Top panel. Heatmap plot catalogues encoded protein locations, ORF amino acid positions and *ω* values at 17D-204 specific codon sites in YFV virulent strains (n=272). Individual heatmap cells present amino acid consensus residues colour-coded according to *ω* values, relative to the scale bar. * indicates significant evidence for purifying selection at the associated codon (Bayesian posterior probability >0.9). Bottom panels. Plot highlights the amino acid residues present at attenuating sites in either the parental AS27 strain (upper) or in 17D-204 strain (lower), and their frequencies in circulating virulent YFV strains (n=272). (*B*) YFV E heterodimer structural predictions highlight vaccine mutations cluster in E-DIII. Top panel. YFV head-to-toe E heterodimers with E-DI, E-DII, and E-DIII highlighted in red, yellow and blue, respectively. The fusion peptide is highlighted in orange, while membrane proximal and transmembrane helices are coloured grey. Structures are presented in ribbon format with individual 17D-204 specific-mutations highlighted in space-filling format. Bottom panel. E heterodimer surface representations with 17D-204 attenuating mutations coloured pink. (*C*) N-linked glycosylation in E. Upper panel. Sequence logo highlights frequency of N-linked glycosylation motifs in virulent YFV strains (NXS/T) at E residues 269-271, with flanking residues, (n=272). Lower panel. All strains used in this study lack an N-linked glycosylation site at E residues 269-271. (*D*) Heatmap visualizes intrinsic control gene and YFV-receptor mRNA expression from immunodeficient Huh7.5.1 hepatoma cells, as determined by RNA-seq. TPM: Transcripts per million mapped reads. (*E*) Virulent YFV and 17D-204 enter Huh7.5.1 cells by CME. Infections were performed with the indicated strains (MOI 1) in the presence of CME inhibitor Pitstop2 or DMSO control, and vRNAs determined at 24 hpi. Data points represent values normalized to DMSO treated controls (100%) from n=3 biological replicates ± SEM. **** *p*<0.0001. Hpi: hours post infection. (*F*) Time-course comparison of virion secretion from Huh7.5.1 infected with the indicated YFV strains (MOI 0.01). Data represent mean PFUs from n=3 biological replicates ± SEM. * *p*<0.05, ** *p*<0.01, ns = non-significant. PFU: plaque forming units. (*G*) 17D-204 results in enhanced apoptotic cell death compared to virulent strain infections in Huh7.5.1 cells (MOI 0.01). Cell viability was determined by flow cytometry at 72 hpi using ApoTracker dye and 7-AAD. Data points represent values from n=3 biological replicates ± SEM. ** *p*<0.01, **** *p*<0.0001, ns = non-significant.

### Vaccine-specific mutations from a surface exposed patch on E-domain III

Vaccine-specific mutations were distributed throughout the YFV ORF with over-representation in the E protein (12 of 30 sites) [38]. E protein heterodimers decorate the surface of mature YF virions, mediating entry into permissive cells. Previous studies reported improved cell uptake for vaccine compared to virulent YFV, associated with E specific mutations [24,25]. To further explore E-mediated entry associated with 17D-204, we first investigated the spatial organisation of vaccine-specific mutations in E by generating structural predictions using Alphafold [39]. YFV E is a prototypic class II fusion protein and predicted structures mirrored experimental structures with high confidence, with recapitulation of three defined β-sheet-rich domains (E-DI, E-DII, and E-DIII) and heterodimer organization in the prefusion conformation [40] (Supplementary Fig. S1). Alphafold modelling also allowed us to visualize membrane proximal α-helices and transmembrane domains, which are absent from experimental structures. Mapping attenuating mutations onto prefusion YFV E heterodimers confirms their spatial arrangement, with mutations absent in transmembrane domains and the fusion peptide (Fig. 2b upper panel). These analyses confirm a surface exposed cluster of YFV 17D-204 specific mutations observed on E-DIII, which overlaps the YFV receptor binding domain (RBD) in the pre-fusion conformation.

### Vaccine and virulent strains used possess identical E glycosylation patterns

The original 17D seed lot no longer exists, but two distinct vaccine sub strains are in use globally: 17D-204 and 17DD [41]. Previous reports have noted differences in N-linked glycosylation between vaccine sub-strains. Notably, 17DD contains an N-linked sequon at E residues 269-271 which is absent in 17D-204 [42]. Only <u>N</u>X<u>S/T</u> amino acid sequons have sugar moieties attached. We investigated the prevalence of N-linked sequons in globally sampled virulent YFV and noted most isolates possess an <u>N</u>N<u>S</u> sequon, although a significant minority did not (Fig. 2c, upper panel). To maximize YFV virulent strain diversity in downstream infection experiments, we compared the 17D-204 vaccine strain with two genetically divergent pathogenic strains: AS27 (WAL II) and UG48 (EAL) (see Fig. 1c). Metagenomic sequencing recovered full-length viral genomes, which were translated into amino acids and compared to 17D-204 (Supplementary Fig. S2). Of note, all strains used here in downstream infection experiments possessed NDN amino acid triplets at residues 269-271: no N-linked glycans are attached to this triplet (Fig. 2c, lower panel). A second <u>N</u>M<u>T</u> sequon at E residues 470-472 was completely conserved among all three strains, and all database derived genomes. Thus, all strains used in this study possess equivalent N-linked glycosylation patterns in E, with differences between attenuated and virulent strains concentrated in E-DIII.

### Vaccine enhanced propagation and apoptosis in hepatoma cells

Huh7.5.1 cells represent human hepatoma cells which are deficient in their ability to respond to viral infection due to impaired RIG-I recognition of viral replicative intermediates, facilitating enhanced viral propagation [43,44]. We reasoned this hepatocyte-derived cell-line would allow us to compare attenuated and virulent YFV cell-uptake and propagation in the absence of the confounding effects of innate immunity. Recently, multiple lipoprotein receptors have been described to mediate YFV entry to cells, including LRP1, LRP4, LRP8 and VLDLR [45,46]. RNA-seq confirmed high mRNA expression of hepatocyte markers and robust expression of all recently described YFV receptors in Huh7.5.1 cells (Fig. 2d). Flavivirus cell-entry occurs via clathrin-mediated endocytosis (CME), with low pH conditions in endosomes initiating gross conformational changes in E which prime virus-host membrane fusion [1,47]. To investigate potential differences in CME-mediated uptake between strains, authentic YFV infections were performed in the presence of the CME inhibitor PitStop2 (Fig. 2e). In contrast to previous reports [25], all strains were highly susceptible to PitStop2 pretreatment (>95% inhibition), indicating both vaccine and virulent YFV enter human hepatoma cells via CME. We next infected Huh7.5.1 cells to monitor potential strain-specific differences in virus propagation (Fig 2f). Infections were performed at a low MOI as differences between strains were masked at higher MOIs. 17D-204 propagation was significantly enhanced at 24 hpi, with 4-8-fold more virions secreted when compared to UG48 and AS27, respectively. Higher 17D-204 production compared to virulent strains, although non-significant, was also seen at 48 hpi but this trend was less visible at 72 hpi (Fig 2f). These data confirm that the 17D-204 strain exhibits an early growth advantage over virulent YFV strains in immune deficient hepatoma cells. Indeed, a cell spreading assay, comparing spreading of the vaccine to its parental strain, confirm enhanced spreading of 17D despite lower MOI infections (Supplementary Fig. S3). YFV represents a cytolytic virus and hepatocyte apoptosis is reported in autopsies from fatal cases [11]. We next investigated differences in the detection of apoptosis markers in vaccine or virulent strain infected Huh7.5.1 cells using a flow-cytometry based assay (Fig. 2g). These analyses highlight significantly enhanced detection of late apoptosis markers for 17D-204 infected cells, when compared to virulent strains at 72 hpi. Together these data confirm that while all YFV strains enter hepatoma cells by CME, the vaccine strain displays an early propagation advantage and is a more potent inducer of apoptosis when compared to virulent strains in innate immune deficient Huh7.5.1 cells.

### Robust 17D-204 vaccine propagation in PHH

YFV primarily targets the liver, although controlled infection studies comparing authentic virulent and attenuated YFV infections in physiologically relevant target cells are currently lacking. To investigate potential differences in hepatotropism and host responses between virulent and attenuated YFV, plated PHH from three donors were infected with either 17D-204, AS27 or UG48 at equivalent MOI (0.5), or left uninfected (mock). Intracellular viral RNA (vRNA) replication and virion production time-courses (24 - 72 hpi) were monitored by RT-qPCR and plaque assay, respectively (Fig. 3a). PHH possess intact innate immunity and respond robustly to viral infections [31–33]. Of note, despite 50 times more viral inoculum (MOI 0.01 vs MOI 0.5), viral titres were 1 log reduced in PHH at 24 hpi when compared to Huh7.5.1 cells (Fig. 2f and Fig. 3a, right panel). For all donors, 17D-204 infections resulted in early peak virus production at 24 hpi, approximately 1 log higher than virulent strains, which slowly declined over the infection time course. Contrastingly, production of genetically divergent virulent strains was initially lower but gradually increased over the infection time course. Similarly, vRNA accumulation profiles were virtually identical for divergent pathogenic strains and distinct for 17D-204. However, differences between vaccine and virulent YFV replication/propagation were generally non-significant. These data highlight temporally structured YFV replication and production kinetics in PHH, which differ subtly between vaccine and virulent strain infections. Similar to Huh7.5.1 infections, 17D-204 possesses a growth advantage at earlier time points in PHH. In summary, these data confirm that the 17D-204 vaccine is highly hepatotropic, propagating robustly in explanted PHH and with earlier peak propagation compared to virulent strains.

**Fig. 3.**
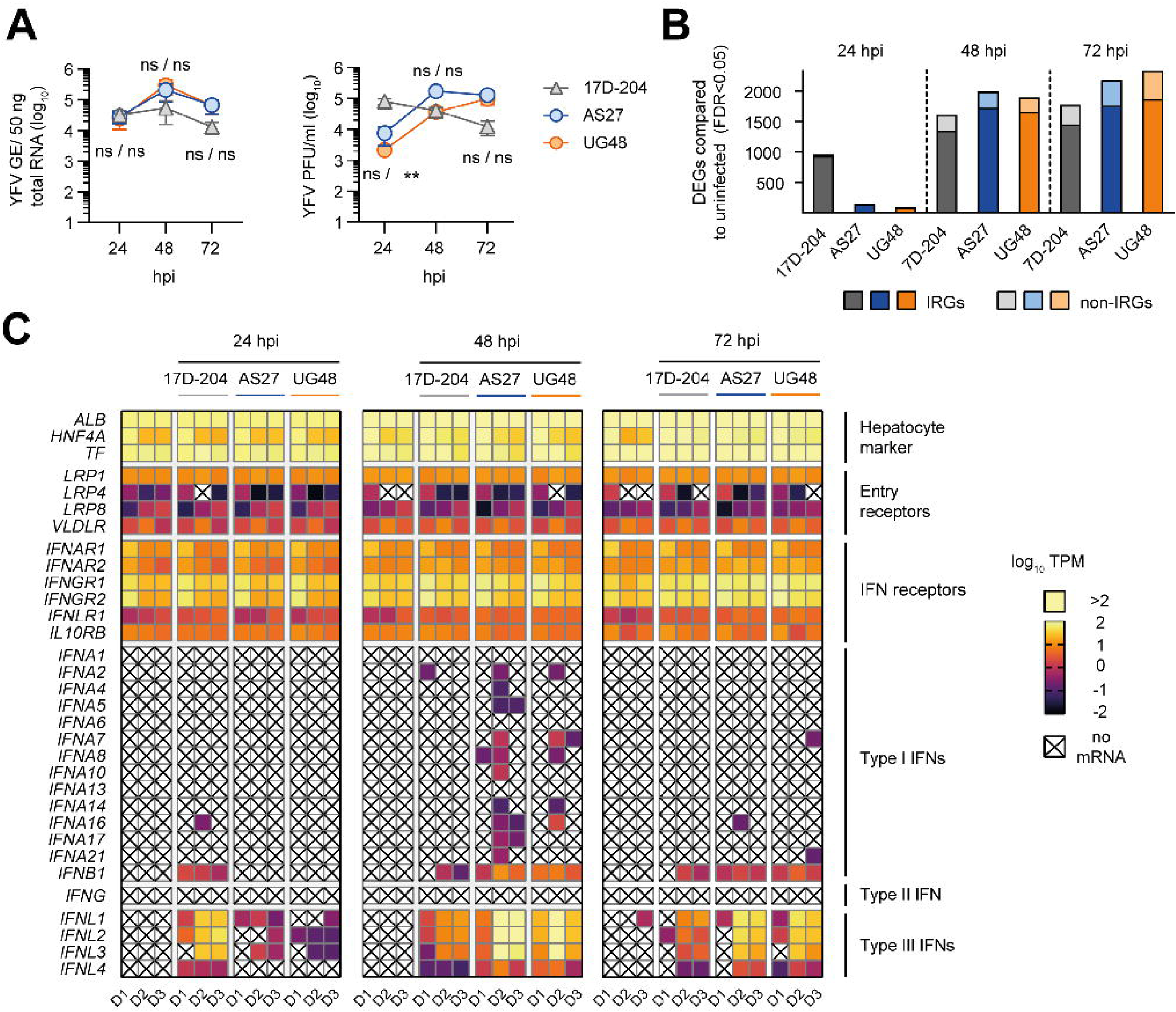
Temporally distinct YFV propagation and IFN induction profiles in PHH. (*A*) Time-course comparison of intracellular vRNA accumulation and virion secretion from PHHs infected with the indicated YFV strains. Data represent mean vRNAs (upper panel) and PFUs (lower panel) from n=3 biological replicates ± SEM. GE: genome equivalents; PFU: plaque forming units; MOI: multiplicity of infection. ** *p*<0.01. ns – not significant. (*B*) Numbers of YFV strain-specific DEGs in PHH at the indicated timepoints, compared to timepoint matched uninfected PHH (n=3 donors). IRGs were determined using Interferome v2.01 [90]. DEG: differentially expressed genes; IRG: IFN regulated gene; FDR: false discovery rate. (*C*) Heatmap visualizes normalized expression of selected mRNA transcripts in PHH infected with the indicated YFV strains, or left uninfected, as determined by RNA-seq. Left panel. 24 hpi. Middle panel. 48 hpi. Right panel. 72hpi. TPM: transcripts per million mapped reads.

### Temporally distinct gene induction profiles for vaccine and virulent YFV in PHH

To compare global transcriptional responses to attenuated and virulent YFV strains, RNA-seq was performed on infected PHH and compared to timepoint matched uninfected controls. RNA-seq confirmed a temporally structured reprogramming of the PHH transcriptional landscape upon YFV infection. Statistical analyses of gene dysregulation in infected PHH quantified differentially expressed genes (DEGs) induced by YFV infection, with distinct profiles observed for vaccine versus virulent strains (Fig. 3b). In line with the dynamics of virion production, broad host transcriptional responses to infection were activated earlier for 17D-204, with 961 DEGs at 24 hpi (FDR p<0.05), a further increase at 48 hpi (1612 DEGs) and maintenance at 72 hpi (1776 DEGs). In contrast, global hepatocyte responses were delayed for virulent strains, with limited activation at 24 hpi (100-151 DEGs), followed by massive amplification at 48 hpi (1896-1994 DEGs), and maintenance at 72 hpi (2183-2335 DEGs) (Fig. 3b). For both 17D-204 and virulent strains at all time points, PHH DEGs were dominated by IFN-regulated genes (80-96%) (IRGs) (Fig. 3b), with the majority upregulated (Supplementary Fig. S4).

### Vaccine and virulent YFV infections induce *IFNB* and *IFNL1-4* in PHH

We next focused on intrinsic messenger RNA (mRNA) expression of hepatocyte markers, YFV receptors, IFN receptors and type I-III IFNs across all experimental conditions (Fig. 3c). In line with expectations, we noted high mRNA expression of hepatocyte markers and detectable mRNA expression of IFN receptors in all samples. In plated PHH, *LRP1* expression was robust and stable across all donors and conditions. In contrast, *VLDLR* expression was generally lower, while *LRP4* and *LRP8* expression was low or completely absent and exhibited more donor specific variability. Mirroring viral propagation kinetics, 17D-204 infected PHH displayed earlier significant induction of *IFNB* and *IFNL1-4* at 24 hpi which was maintained at 48 hpi and declined at 72 hpi. In contrast, PHHs infected with virulent YFV strains displayed delayed induction profiles: limited expression of *IFNB* and *IFNL1-4* transcripts at 24 hpi, followed by significant induction at 48 hpi and maintenance at 72 hpi. Of note, despite *IFNAR* and *IFNGR* receptor expression, no meaningful induction of either *IFNA* subtypes or *IFNG* was detected in YFV infected PHH. In summary, both attenuated and virulent YFV induce expression of *IFNB* and *IFNL1-4* in PHH, with earlier induction observed for the vaccine. We observed robust propagation of all YFV strains in PHH despite broad activation of antiviral responses mediated by *IFNB* and *IFNL1-4*, although considerably reduced when compared to immune deficient Huh7.5.1 cells. Together, these experiments catalogue induction of specific IFNs and concomitant IFN-mediated defence programs in PHH, which are comparable between vaccine and virulent YFV. Our data highlight subtle temporal differences in infection induced cell-intrinsic responses observed between strains, which mirrors earlier propagation by the vaccine in PHH.

### Vaccine and virulent YFV infections induce overlapping gene programs in PHH

To investigate whether distinct cell-intrinsic processes were modulated by attenuated and virulent YFV infections, we performed Gene Ontology (GO) enrichment analyses of infection-induced DEGs, comparing strain-specific biological processes targeted at different time-points. To provide a visual overview of all biological processes targeted, redundancy and repetition in GO enrichment results were reduced by generating semantic similarity matrices, clustering GO terms into related groups at 24, 48 and 72 hpi (Fig. 4a-c). In line with virus propagation profiles and RNA-seq analyses (Fig. 3), significantly enriched GO categories induced by 17D-204 infections were numerically higher than for virulent strains at 24 hpi (Fig. 4a), with more overlap in targeted biological processes observed at 48 and 72 hpi (Fig. 4b and 4c). At 24 and 48 hpi, biological processes associated with virus-induced innate immunity and response to cytokines were numerically dominant in significantly enriched GO terms. While these were still apparent at 72 hpi, an expansion of significantly enriched GO terms associated with metabolism and biosynthesis was observed.

**Fig. 4.**
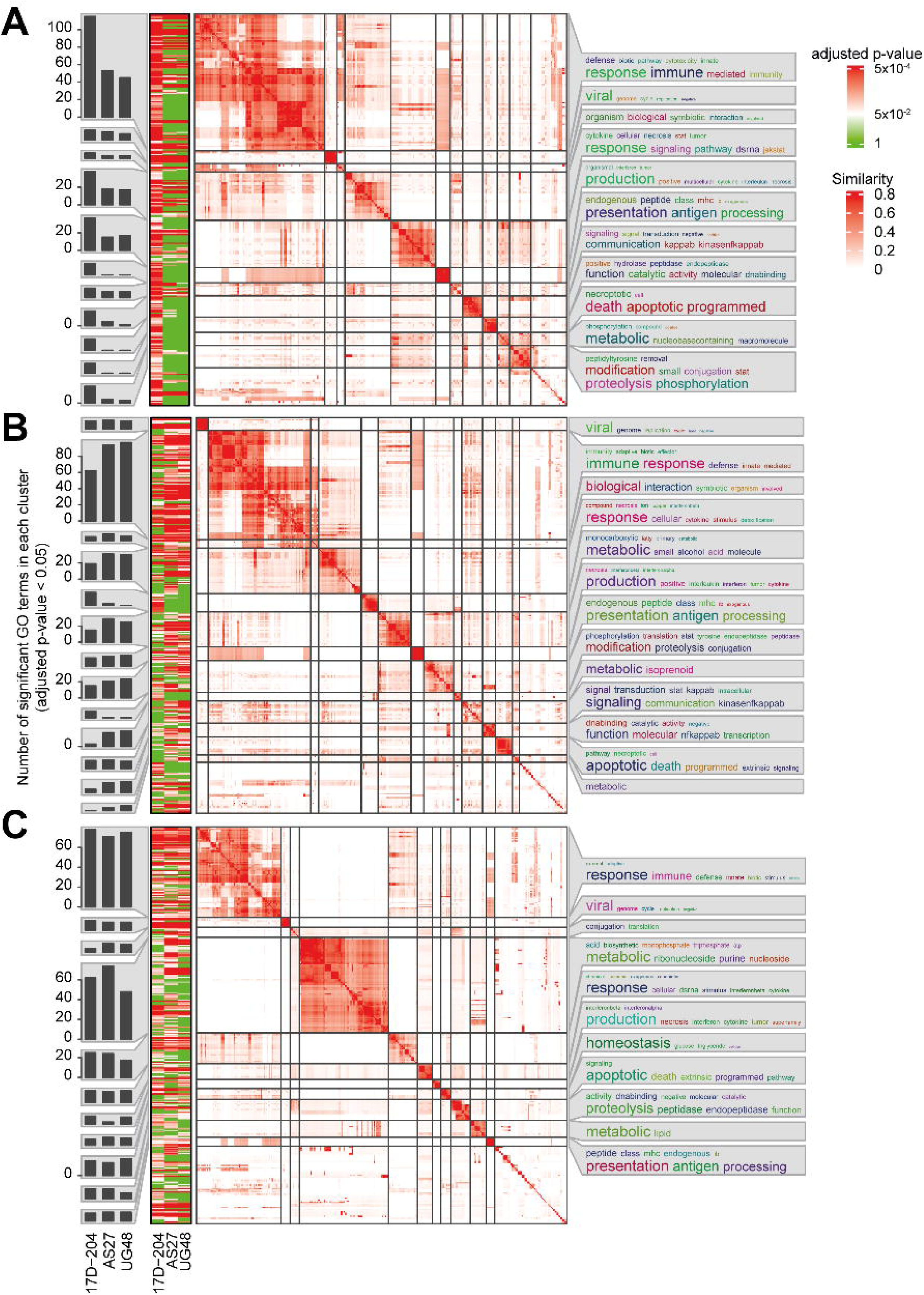
Overlapping YFV infection-induced gene programs in PHH. Heatmaps visualize enrichment of biological process GO terms induced by 17D-204, AS27 and UG48 infections of PHH, with clustering of terms based on semantic similarity. (*A*) 24 hpi (*B*), 48 hpi and (*C*) 72 hpi. Bar plots to the left of the enrichment heatmaps depict the numbers of significantly enriched GO terms in each cluster (FDR *p*<0.05). Word cloud annotation to the right of the semantic similarity heatmap summarizes the biological processes represented in each cluster, with keywords extracted from GO term descriptions. Font size of keywords is proportional to significance. Clusters containing fewer than 10% of the total number of terms are combined into a single cluster at the bottom of each semantic similarity heatmap. FDR, false discovery rate.

Visualization of selected representative GO terms confirms that YFV-induced transcriptional dysregulation targets a broad range of cell-intrinsic processes, with significant modulation gene programs associated with antiviral responses, immunity, cell death, inflammation, metabolism and translation (Fig. 5a). In line with 17D’s earlier propagation in Huh7.5.1 cells (Fig. 2f), coupled with earlier peak propagation and IFN induction in PHH (Fig. 3a and 3c), we observed temporally distinct transcriptional response waves mediated by attenuated and virulent yellow fever virus strains. Overall, dysregulated genes and cellular processes were shared between YFV strains, with earlier activation observed for 17D and delayed but enhanced activation for virulent strains (Fig. 6a, 6b and Supplementary Fig. S5). To further dissect PHH responses to individual YFV strains, we visualized induction profiles of selected subsets of transcripts associated with distinct cellular processes affected by infection, including those encoding RNA sensors, antiviral transcription factors (TFs), antiviral effectors, human leucocyte antigens (HLAs), caspases and chemotactic chemokines/cytokines, metabolic genes and ribosomal proteins (Fig. 6c and Supplementary Fig. S6). In-line with DEG and GO analyses, these data reveal that infection of PHHs with attenuated or virulent YFV modulates overlapping expression of virtually identical transcripts. Consistent with virus propagation and IFN induction profiles, these data highlight differences in the temporal dynamics of immune activation between vaccine and virulent strains and confirm that dysregulated cellular gene programs are dominated by host immunity, response to cytokines and metabolic reprogramming in PHH. Irrespective of the infecting strain, these data also highlight propagation of both vaccine and virulent YFV in PHH despite activation of type I and III IFNs and concomitant antiviral responses.

**Fig. 5.**
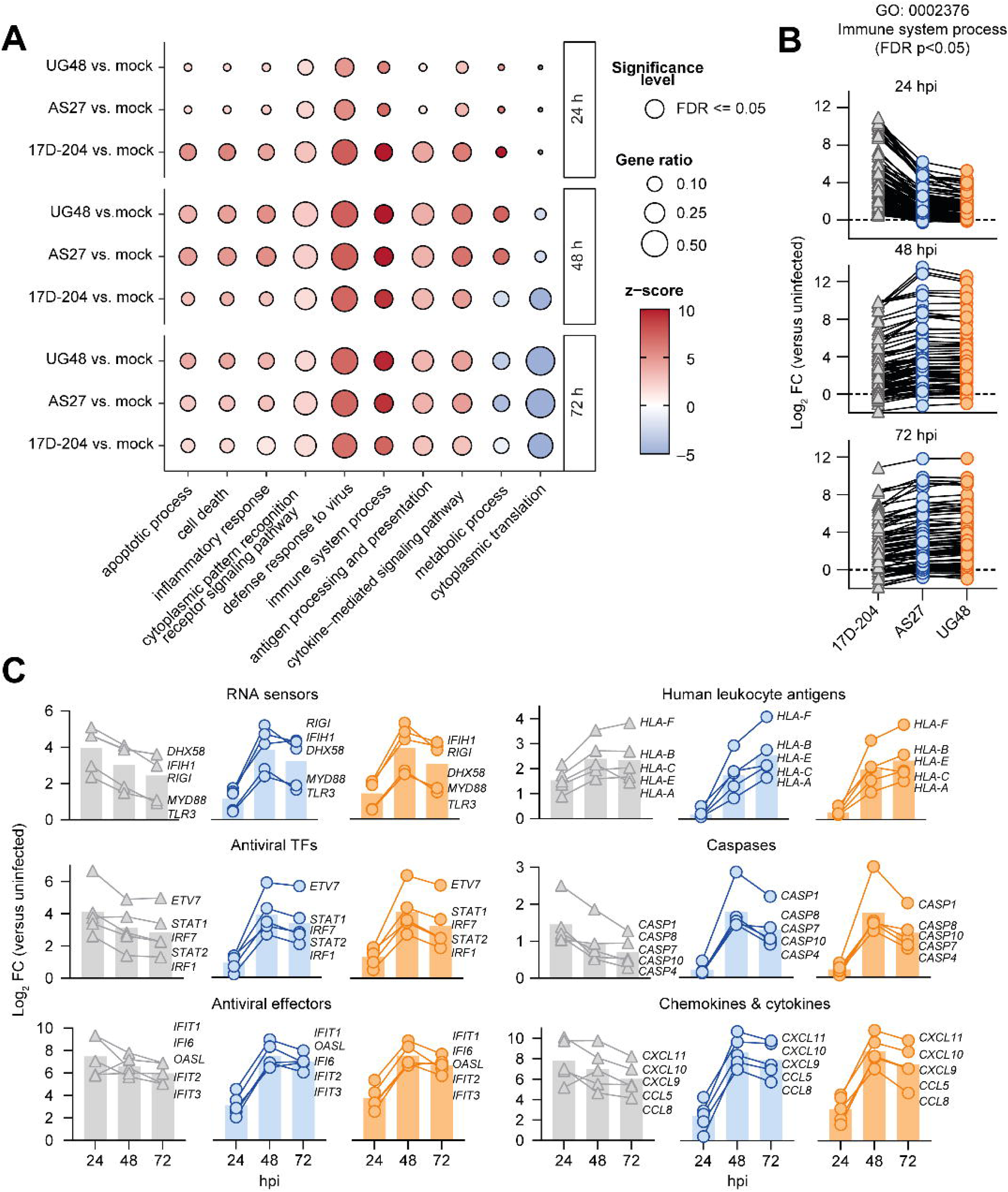
Vaccine and virulent YFV induce comparable gene programs in PHH. (*A*) Enriched GO terms in YFV-infected PHHs encompass diverse biological processes and are temporally structured. Dot-plot visualizes enriched GO categories which are shared between strains across sampling points but exhibit temporally distinct induction kinetics. (10 representative categories shown). (*B*) Magnitude of induction for individual genes in the GO category ‘Immune System Process’ differs between strains across time points. Genes were selected based on significant differential expression when comparing vaccine and virulent strains (FDR <0.05). Plots compare induction of identical genes at distinct timepoints. Individual data points represent induction (log2 fold change) compared to timepoint matched uninfected PHH and represent the mean induction for n=3 donors infected with the indicated YFV strains. (*C*) YFV strain-specific gene profiles induction in PHH. Plots compare induction profiles for selected categories of immunity genes encoding RNA sensors, kinases, transcription TFs, antiviral effectors, HLAs, caspases and inflammatory chemokines/cytokines (n=5 genes per category). Individual data points visualize gene induction (log2 fold change) compared to timepoint matched uninfected PHHs and represent the mean induction for n=3 donors infected with the indicated YFV strains. TF: transcription factor

**Fig. 6.**
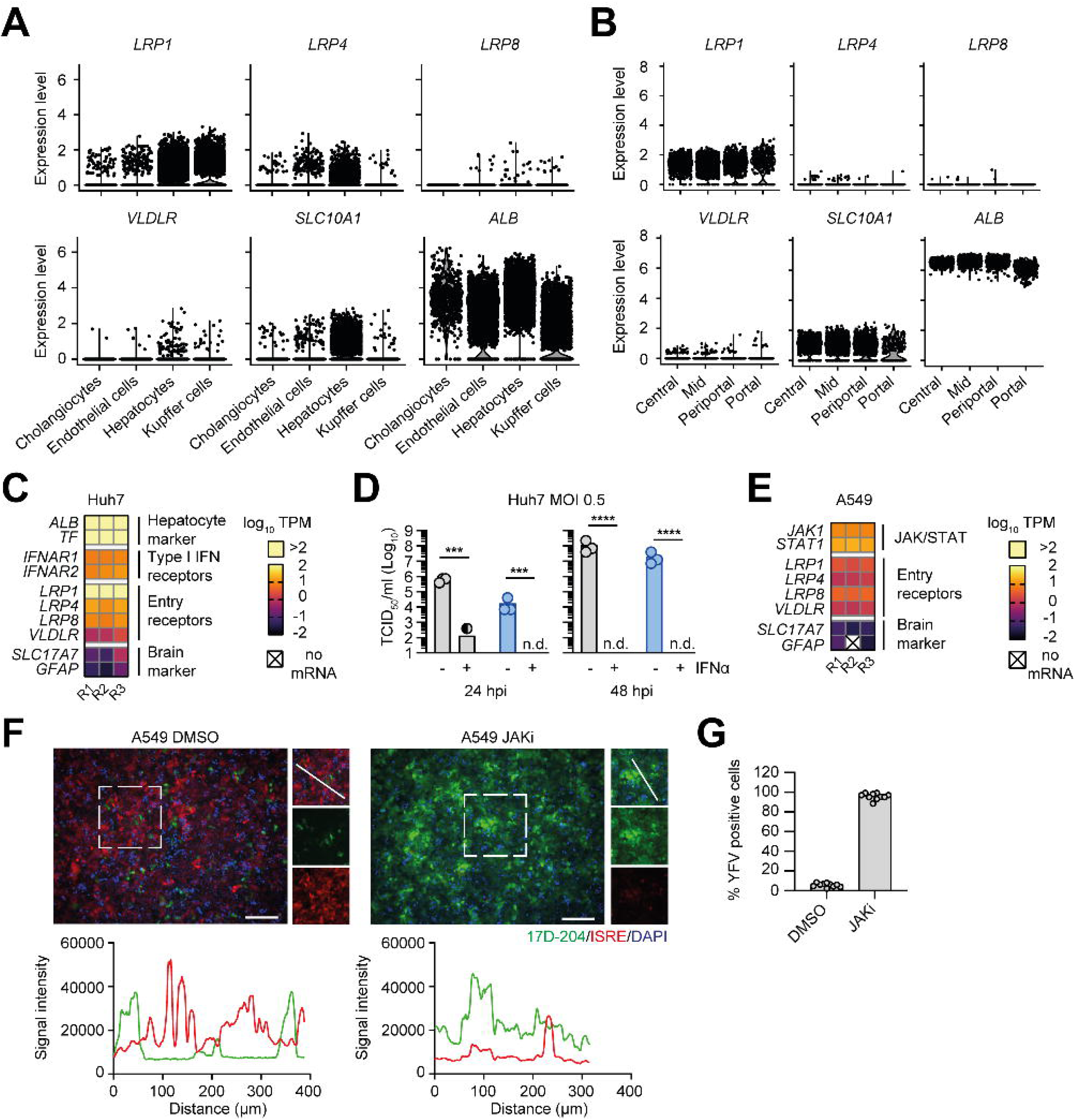
IFNα and paracrine signalling restrict YFV spread. (*A*) YFV receptor transcript expression across liver resident cell-types. Plots represent normalized gene expression for multiple YFV receptors (*LRP1*, *LRP4*, *LRP8* and *VLDLR*), the HBV receptor (*SLC10A1*) and control liver marker (*ALB*), as determined by scRNA-seq. (*B*) YFV receptor gene expression across defined liver zones, determined by Visium spatial transcriptomics. For (*A*) and (*B*), data was retrieved from https://www.livercellatlas.org/. (*C*) Heatmap visualizes intrinsic control gene and YFV-receptor mRNA expression from Huh7 cells, as determined by RNA-seq. TPM: Transcripts per million mapped reads. (*D*) Attenuated and virulent YFV propagation is abrogated by IFNα pretreatment. Infections were performed in Huh7 cells with either 17D-204 (grey) or AS27 (blue), with or without IFNα pretreatment. Viruses secreted into supernatants were determined by TCID50 at 24 and 48 hpi. Data points represent values from n=3 biological replicates ± SEM. Hpi: hours post infection. *** *p*<0.001, **** *p*<0.0001. n.d.: none detected. (*E*) Heatmap visualizes intrinsic control gene and YFV-receptor mRNA expression in A549 cells, as determined by RNA-seq. (*F*) YFV 17DvGFP infection of A549 ISRE-mCherry cells (MOI 10, 48 hpi). Immunofluorescence images visualize mock treated, YFV infected cells (left panel) and baricitinib pretreated (JAKi), YFV infected cells (right panel). Insets highlight specific regions used for quantification of 17D-204 (green) and ISRE-mCherry (red), with signal intensity plots positioned directly below. (*G*) Quantification of YFV foci in A549 ISRE-mCherry cells with DMSO or baricitinib (JAKi) treatment.

### Enriched *LRP1* expression in the liver

YFV is the only member of the Orthoflavivirus genus that consistently causes severe liver pathology [27]. To investigate determinants which contribute to YFV’s unique hepatotropism, we first leveraged publicly available scRNA-seq and Visium data sets available through Liver Atlas (https://www.livercellatlas.org/) to visualize recently discovered YFV receptor transcript abundance in the liver at single-cell and spatial resolution, which has not been previously described. As control genes, we included hepatocyte-specific *SLC10A1* (encoding NTCP, the hepatitis B virus receptor) and *ALB*, which is highly expressed in the liver. Our bulk RNA-seq data highlights robust *LRP1* receptor expression in PHH across all conditions and donors (Fig 3c). The scRNA-seq dataset highlights robust *LRP1* expression in most hepatocytes and Kupffer cells, with fewer cholangiocytes and endothelial cells expressing this gene (Fig. 6a). *LRP4* expression was also enriched in hepatocytes, while *LRP8* and *VLDLR* expression was detectable across all cell-types but detected in comparatively few cells. Liver zonation is the functional specialization of hepatocytes in different zones of the liver lobule. Consistent with scRNA-seq data, the Visium spatial transcriptomics dataset from liver biopsies confirmed abundant *LRP1* expression across the central, mid, periportal and portal zones (Fig. 6b). In contrast *LRP4*, *LRP8* and *VLDLR* expression was detectable, but restricted to relatively few cells. Together, these datasets allow determination of YFV receptor expression in the major target organ at unprecedented resolution and confirm *LRP1* as the dominant receptor likely used by YFV to infect the liver. Of note, differences were observed between scRNA-seq and Visium data concerning *LRP4* expression. These data also highlight the potential for parallel YFV entry routes into individual cells, with a small population of liver resident cells expressing multiple receptors simultaneously.

### IFNα Inhibits YFV propagation in Huh7 cells

Despite robust *IFNB* and *IFNL1-4* induction, we observed no meaningful induction of *IFNA* subtypes in PHH upon infection with YFV (Fig 3c). To further investigate determinants which contribute to YFV’s hepatopathology, we tested inhibition of YFV propagation by IFNα pretreatment in human hepatoma cells. For these experiments we utilized Huh7 cells which exhibit immune reactivity to YFV infection [24] and express *IFNA* receptor mRNAs plus all YFV entry receptors (Fig. 6c). Cells were infected with YFV, with or without IFNα pretreatment, and secreted virions were quantified at 24 and 48 hpi (Fig. 6d). These data confirmed that YFV propagation was almost completely inhibited by IFNα pretreatment, highlighting susceptibility of both vaccine and virulent YFV to IFNα inhibition.

### Paracrine IFN signalling restricts YFV 17D-204 spread in A549 cells

From the bulk RNA-seq data generated, we quantified global PHH responses to YFV infection (Fig. 3-5), although whether these responses are dominated by autocrine or paracrine IFN signalling remained unclear. Due to technical limitations associated with primary cells and limited donor material, we were unable to evaluate this question directly in PHH. To provide insights, we utilized A549 cells which react strongly to viral infections and immune stimulus with robust type I and III IFN responses [24,43,48], and express all YFV receptors (Fig. 6e). We combined this with a YFV-17D venus GFP (vGFP) reporter virus [49] which allowed us to visualize individual virus-infected cells. To facilitate visualization of IFN-mediated innate immune activation in individual cells, A549 cells were transduced with an IFN-stimulation response element (ISRE) mCherry reporter construct. Infection of A549-ISRE-mCherry cells with YFV-17DvGFP virus highlights minimal overlap between YFV infected cells and activated ISREs, highlighting the dominant role of paracrine signalling in the bulk cellular response to YFV infection in these cells (Fig. 6f). This effect could be ablated by baricitinib treatment, which blocks JAK/STAT signalling, ablating IFN production and enhancing virus infection rates by approximately 90% (Fig. 6f and 6g). In summary, IFN-mediated cellular responses to infection were dominated by paracrine signalling in YFV-infected A549 cells, presumably due to viral mediated suppression of autocrine responses in infected cells and paracrine activation in bystander cells.

## Discussion

In our study, we first utilized database deposited sequences to elucidate drivers of YFV genome diversification. The evolutionary origins of YFV lie in tropical Africa, with import of the virus to the South American continent associated with European colonialism and the transatlantic slave trade [34–36]. Employing phylogenetic methods, previous studies have reported the evolutionary relationships of both African and globally sampled YFV strains [34,36,50]. However, YFV represents a neglected tropical disease [51]. Consequently, due to historically sparse genomic sampling of YFV in endemic regions coupled with the size limitations associated with Sanger sequencing approaches, previous evolutionary analyses were performed using subgenomic fragments often encoding single proteins (e.g. prM) and derived from limited numbers of viral isolates. More recently, the advent of metagenomic sequencing approaches has revolutionized pathogen surveillance, facilitating the genomic monitoring of YFV outbreaks in real-time [52]. Leveraging this recent accumulation of database deposited full-length or near full-length YFV genomes, we investigated the phylogenetic relationships and evolutionary processes which shape global virulent YFV diversity at the genomic level, before focusing on individual codon-sites where vaccine attenuating mutations occur. We implemented site-by-site codon analysis, scanning aligned YFV ORFs from nearly 300 globally sampled genomes from virulent isolates. These analyses highlight purifying (negative) selection as the major evolutionary force acting on YFV protein-coding sequences – the majority of YFV ORF codons exhibit a significant excess of synonymous substitutions when compared to non-synonymous substitutions (Fig. 1d). Focusing on codons at which vaccine-specific mutations occur, we identified significant evidence for purifying selection at 28/30 codons, implying functional constraints associated with effective YFV transmission and propagation limit amino acid residue variability (Fig. 2a). Indeed, multiple specific selective constraints are imposed by the *Orthoflavivirus* transmission cycle: YFV experiences sequential bottlenecks upon continued cycling through mosquito vectors, coupled with requirements for host factor compatibility in both vertebrates and arthropods which diverged ∼700 million years ago [53]. Vaccine-specific mutations confer deleterious phenotypes to virulent YFV by dramatically reducing systemic viremia in infected humans and nonhuman primates [41] and abrogating mosquito transmission due to midgut barriers which prevent systemic dissemination [54]. The serial passage of 17D in mouse brains, and subsequently minced chicken embryos with the brain and spinal column removed, eliminated selection pressures associated with propagation in mosquitos and systemic visceral infection of primates [15,16]. However, chicken embryos are innate immune competent and can actively recognize and respond to viral replication [55,56]. Thus, accumulation of attenuating mutations during YFV serial passage results from removal of selective constraints associated with the natural viral transmission cycle and systemic dissemination, rather than an absence of innate immune targeting. Together, this highlights an overlap between the viral determinants required for successful YFV replication and dissemination in mosquito vectors, and viscerotropic dissemination, high systemic viremia and associated pathology in primates. Indeed, the signatures of strong purifying selection within codons where vaccine-specific mutations occur confirms their importance in the natural YFV transmission cycle, as demonstrated by the extreme amino residue conservation at these positions in genetically divergent virulent strains.

Over-representation of vaccine attenuating mutations is observed in E, which is responsible for cell-entry, elicits protective neutralizing antibody responses and is the YFV protein exhibiting the highest level of amino acid conservation (Fig. 1c). Visualization of vaccine-specific E mutations in a structural context identifies residues localized on the heterodimer surface. Surface exposed sites were located in E-DI and E-DII, with a notable cluster of four vaccine-specific mutations located in E-DIII forming a discrete surface exposed patch (Fig. 2b). Pre-fusion E-DIII on the virion surface forms the first YFV-host interface by direct interaction with host plasma membrane localized proteins, engaging directly with host receptors to initiate infection [45,46,57,58]. The concentration of attenuating mutations in YFV E-DIII likely modulates 17D cell-tropism, enhancing early propagation and cell-spreading (Fig. 2f, Supplementary Fig. S3, and [24]). Additionally, we noted increased cytolytic responses induced by 17D compared to virulent strains. These data reflect observations from our previous studies on additional viruses, where more efficient cell-entry is associated with earlier innate immune induction and more pronounced cell death for SARS-CoV-2 [43], and adaptive mutations which enhance replication and cell-to-cell spreading in tissue culture for HCV, but result in stronger innate immune induction and mild attenuation in PHH [33]. Apoptosis helps to control the magnitude and duration of immune responses, preventing overstimulation and excessive inflammation. However, hepatocyte apoptosis is a major histopathological feature observed in virulent YFV liver pathology [11,27,28] suggesting this property is not a correlate of 17D’s attenuation.

Virulent YFV infection causes systemic viremia and viscerotropic disease but primarily infects the liver, which is the site of the most significant tissue damage [11,27]. Here, the virus is detected early in Kupffer cells and later in hepatocytes, where it impairs excretory, synthetic and metabolic functions, causing jaundice and leading to hepatocyte apoptosis (Councilman bodies) and midzonal necrosis [11,27,28,50]. By contrast, vaccination in humans and nonhuman primates is associated with localized infections, transient low-level viremia and an absence of associated liver damage [15,41,59,60]. These data imply that during attenuation 17D may have either: (a) lost the ability to infect hepatocytes, (b) acquired impaired immune innate antagonism compared to virulent YFV resulting in rapid hepatocyte clearance or, (c) retained the ability to infect hepatocytes and cause liver pathology, which is normally prevented by impaired dissemination to the liver. While previous studies have compared vaccine and virulent strain propagation in hepatoma cells [24,61], the implications of robust vaccine replication in human liver cells were previously overlooked, specifically with respect to the absence of liver pathology upon vaccination. Indeed, whether vaccine strain infection of hepatocytes occurs after immunization with 17D in healthy individuals remains unclear. A comprehensive Brazilian vaccination cohort study provided compelling evidence for impaired systemic dissemination of 17D in humans, with limited incidences of systemic viremia or elevated liver enzymes in vaccinees [59]. Our data highlight that in the absence of potential physiological barriers associated with systemic *in vivo* infections, the YFV 17D vaccine can productively infect PHH *ex vivo*. We observed comparable propagation and near-identical host transcriptional responses to vaccine and virulent YFV strains in PHH, although subtly different temporal kinetics were observed (Figs. 3-5).

Infection with 17D was previously shown to induce metabolic perturbations including mitochondrial hyperactivity, which enhances innate immune activation [62]. Furthermore, 17D infected cells exhibit elevated expression of selected pro- and anti-inflammatory cytokines when compared to Asibi infected cells [63], and also binds to cells more efficiently than Asibi, resulting in increased cytosolic delivery of vRNA which mediates enhanced production of selected cytokines [25]. The E and NS2A proteins were reported as key enhancers of vaccine spread and concomitant antiviral responses [24]. Together, these studies report amplified innate immune induction by multiple mechanisms associated with 17D infections. In contrast to these reports, our PHH infection time-course data does not support enhanced innate immune induction by the 17D compared to virulent strains, but rather earlier activation by the vaccine due to earlier peak propagation in PHH (Fig. 3a). Indeed, the breadth and magnitude of PHH antiviral responses to virulent YFV is elevated compared to 17D at later time points (Figs. 3-5, Supplementary Fig. S5), corresponding with delayed peak propagation. Consistent with our previous observations in PHH [32], IFN induction by YFV was also restricted to *IFNB* and *IFNL1-4*. This was despite detectable expression of IFN I-III receptors and indicates that hepatocytes are unable to produce type I IFNα subtypes at meaningful levels, irrespective of the infecting virus. Despite vigorous induction of *IFNB* and *IFNL1-4* mediated defence programs, robust vaccine and virulent YFV propagation occurs in PHH. This is in stark contrast to HCV, where infection-induced innate immunity in PHH suppresses virus replication and completely abrogates virus production, which can be rescued by JAK/STAT inhibition [31]. We propose YFV’s strong hepatotropism and the associated liver damage caused by infection may be partly mediated by an inability of PHH to produce meaningful levels of IFNα. Indeed, propagation of both vaccine and virulent strains in human hepatoma cells was robustly inhibited by IFNα pretreatment (Fig. 6d). Furthermore, spreading of 17D-204 was dramatically enhanced by JAK/STAT inhibition in cells that produce type I and III IFNs (Figs 6f and 6g). We also propose that the transcriptional response signatures observed for 17D-204 in PHH do not represent the signatures of attenuation, as they are virtually identical to those induced by liver pathology causing virulent strains (Figs. 4 and 5), and exhibit overlap with signatures induced by liver pathology causing viruses HCV [31] and SARS-CoV-2 [32]. Indeed, the temporally distinct transcriptional response waves observed for vaccine and virulent YFV likely reflect differences in propagation kinetics rather than signatures of attenuation.

In humans, vaccination does not induce enhanced immunity compared to infection, with vaccine breakthrough infections occurring [20]. Vaccination with 17D causes a subclinical infection and the process of attenuation resulted in a loss of viscerotropism and impaired systemic dissemination in nonhuman primates [15,41]. However, in very rare cases, 17D vaccination causes yellow fever vaccine-associated viscerotropic disease (YEL-AVD), with symptoms and viral dissemination which are indistinguishable from wild-type disease [64]. YEL-AVD is reportedly associated with polymorphisms in *IFNAR1*, *CCR5* and *RANTES* genes in humans [65,66] and the generation of auto-autoantibodies to IFNα2 is also reported in life-threatening adverse reactions to 17D vaccination [67]. Although mice do not faithfully recapitulate YFV disease phenotypes seen in humans and nonhuman primates, 17D dissemination did not occur in immunocompetent mice but became systemic in a proportion of *IFNAR^-/-^* mice and was associated with limited disease [68]. Furthermore, three vaccine-specific mutations in domain III of the 17D E protein were shown to be involved in reduced neurotropism and attenuated virulence in *IFNAR^-/-^ IFNGR^-/-^* mice [69]. Together, these studies highlight functional innate immunity represents an integral component which impairs 17D’s systemic dissemination and may contribute to restricting liver access.

The recent discovery of multiple YFV entry receptors - LRP1, LRP4, LRP8 and VLDLR - allowed us to visualize their mRNA expression in cell-lines, explanted PHH and publicly available scRNA-seq and Visium data from liver tissue. These data confirm LRP1 as the major liver-expressed receptor which likely facilitates the majority of YFV infection in hepatocytes, although no obvious differences in receptor usage were reported between the vaccine and virulent strains [45,46]. Consistent with this observation, we observe comparable hepatotropism by both vaccine and virulent strains, with robust propagation in PHH. The limitations of our *ex vivo* PHH infections do not allow us to determine whether 17D reaches the liver after immunization *in vivo*, although PHH are highly susceptible and permissive. Whether the 17D vaccine productively infects the liver in healthy human vaccinees is challenging to determine. Therefore, to address this and determine whether 17D can induce liver pathology *in vivo*, future studies bypassing potential physiological barriers which could limit liver infection upon vaccination should be performed in permissive animal models. This could be achieved by hydrodynamic injection of 17D directly into the livers of nonhuman primates, hamsters or human liver chimeric mice, which exhibit similar degrees of liver pathology to humans upon infection with virulent YFV strains. [27,28,70–72].

In summary, while the attenuation of 17D is likely to be multimodal *in vivo*, our data indicate it does not correlate with reduced hepatotropism or innate immune antagonism: vaccine and virulent YFV propagate and induce similar transcriptional responses in explanted PHH. We propose that in healthy individuals, unknown barriers which impair systemic dissemination normally block the vaccine from reaching the liver, supported by the general lack liver enzyme elevation in vaccine cohorts [59]. Indeed, our PHH data suggests that in the absence of such barriers [65,68], the 17D-204 strain is not attenuated for infection of hepatocytes, and takes a similar course to virulent YFV, explaining YEL-AVD. More broadly, our data also confirm an impaired ability of human hepatocytes to produce IFNα subtypes upon viral infection, which may also contribute to YFV’s liver tropism. Together our data yield novel insights into 17D’s virulence attenuation and highlights impaired hepatotropism does not contribute to the vaccine’s excellent safety profile.

## Materials and methods

### YFV strains

The YFV 17D-204 vaccine strain (Stamaril^®^: Sanofi Pasteur) was provided as a kind gift from Sandra Ciesek (Institute for Medical Virology, University Hospital, Goethe University, Frankfurt). Pathogenic YFV strains UVE/YFV/1927/GN/Asibi and UVE/YFV/1948/UG/MR896 TVP3236 (AS27 & UG48) were purchased from the European Virus Archive Global (EVAg). EVAg strains were provided as lyophilized stocks, resuspended in ddH2O, aliquoted and stored at −80°C prior to propagation. Reverse genetics YFV constructs were used for IFN sensitivity experiments, cell spreading assays and A549 ISRE-mCherry infections were pACNR-FLYFV-17Da encoding YFV-17D (YFV-17D ic) [73,74] and pYFV-Asibi ic [75], and YFV-17D venus GFP [49] Virus stocks were produced as described previously [76].

### Tissue culture, YFV propagation and titration

BHK-21, Huh7, Huh7.5.1, VeroE6 and A549 cells were maintained in maintenance media (DMEM supplemented with either 5 or 10% FCS, and with or without 100 Units/mL penicillin, 100 μg/mL streptomycin and 2 mM L-glutamine) at 37°C with 5% CO2. For propagation of YFV stocks, BHK-21 cells were seeded at 1 × 10^7^ cells/T175 flask in maintenance media. After 24 h, maintenance media was exchanged for 5% FCS-containing DMEM spiked with individual YFV strains (MOI 0.1) and incubated for 1 h at 37°C. Subsequently, inoculating media was replaced with fresh maintenance media and infected cells were incubated for 4 days. Upon visible cytopathology, supernatants were harvested and centrifuged at 3,000 ×*g* for 10 min to remove cellular debris. Stocks were further concentrated using Vivaspin 20 Centrifugal Concentrators (Sartorius), with a cutoff size of 100kDa, via centrifugation at 3000 ×*g* for 15min to reduce the volume to at least one third of the initial input. Viral titres were determined by plaque assay after one freeze-thaw cycle. All viruses were aliquoted and stored at −80°C until further use in infection experiments.

For plaque assays, VeroE6 cells were seeded one day prior to infection in 6-well plates. Virus containing supernatants were serially diluted (10*^−^*^1^ to 10*^−^*^5^) prior to transfer to seeded VeroE6 cells. Dilutions were incubated on cells for 1 h before replacement with plaque medium (3.5% Avicel PH-101 in PBS and maintenance media). Avicel overlaid cells were then incubated for 5-6 days. On the final day, plaque medium was discarded, and cells were washed carefully with PBS. Cells were fixed with 4% paraformaldehyde (PFA) in PBS for at least 30 min. Post-fixation, PFA was discarded, and cells were incubated with crystal violet staining solution (1% crystal violet in 20% methanol) for 30 min. The staining solution was then removed, and cell monolayers were carefully rinsed with water. Plates were then air-dried overnight before plaques were counted and viral titres calculated. Experiments with 17D-204 were performed under biological safety level 2 (BSL2) conditions while AS27 and UG48 viruses were handled under BSL 3 conditions, in accordance with authorized procedures for the state of Hessen. For all infection experiments in cell-lines or primary cells, virus inoculations were performed for 1 hour at the stated MOI.

### Cell-spreading and IFN sensitivity assays

To investigate susceptibility to inhibition by IFN, Huh7 cells were treated with 1000 IU/ml IFNα (Sigma-Aldrich/Merck: IF007) prior to infection with YFV-17D ic and Asibi ic stocks at an MOI of 0.5. Infection was followed by a second IFN treatment. Tissue culture infectious dose 50 (TCID50) titration was performed to determine secreted viral titers. Supernatants were added to BHK-21 cells in a serial dilution of 10^-1^-10^-11^. The TCID50/ml was calculated using the Reed and Muench calculator at 5 dpi.

### Primary human cells

Cryopreserved PHH were purchased from either Thermo Fisher Scientific or PRIMACYT Cell Culture Technology. All donors tested serologically negative for HBV, HCV, HIV, CMV and SARS-CoV-2 and individual donor information is detailed below.

| Lot# | Pathology | Gender | Race | BMI |
| --- | --- | --- | --- | --- |
| HU8406 | Cerebrovascular Stroke | Female | Guamanian/Chamorro | 20.6 |
| CHM2221 | Cholangiocarcinoma | Male | Caucasian | 24.3 |
| CHM2225 | Hepatocellular carcinoma | Male | Caucasian | 28.0 |

PHHs were thawed and seeded at a density of 2.5×10^5^ live cells/well on collagen-coated 24-well plates one day prior to YFV infection. In line with the provider’s recommendations, plated PHHs were maintained in a 37°C incubator with 5% CO2 in either William’s E Medium (CM4000, ThermoFisher) or Human Hepatocyte Maintenance Medium (HHMM-500, PRIMACYT), with the recommended supplements added to maintenance media immediately prior to use. Liver biopsies were removed during surgery and would otherwise be discarded as biological waste. Informed patient consent was stated on the provider’s website.

### A549 ISRE reporter assay

Plasmid pLenti ISRE-mCherry was generated from the pLentiPGK Puro DEST JNK KTR Clover backbone (Addgene plasmid #59151). ISRE elements were incorporated as previously described [77]. Assembly was performed using Gibson Assembly (NEB, E2611L) with inserts amplified using Q5 High-Fidelity DNA Polymerase (NEB, M0491L) and primers containing the appropriate overhangs. To produce lentiviral particles, 4 × 10^5^ 293T cells were seeded on collagen-coated 6-well plates. The following day, the 293T cells were transfected with the plasmids pcz-VSV-G, pCMV-dR8.74 and pLenti ISRE-mCherry using Lipofectamine 2000 (Invitrogen, Cat.40 Nr. 11668019) following the manufacturer’s instructions. Six hours post transfection, the medium was changed and lentiviral particles were harvested 48 h post transfection. Supernatants were filtered (Filtropur 0.45, Sarstedt, Cat. Nr. 83.1826) and supplemented with HEPES and polybrene and either used directly or stored at −80 °C. For transduction, A549 cells were seeded on a 6-well plate and inoculated with 1 mL of lentiviral particles for 6 – 8 h. Selection of the transduced cells was started 48 h post transduction using 2.5 μg/mL puromycin. A549 ISRE-mCherry reporter cells were seeded into 96-well plates (2 × 10⁴ cells per well). Twenty-four hours later, the culture medium was removed, and cells were infected with YFV-17D venus GFP (MOI 10). After 48 hours cells were fixed with paraformaldehyde (PFA), and nuclei were counterstained with Hoechst prior to imaging.

### Flow Cytometry

To determine expression levels of apoptosis markers, Huh7.5.1 cells were infected with 17D-204, AS27 and UG48 at an MOI of 0.01, or left uninfected. At 72 hpi, cells were trypsinized, pelleted and re-suspended in PBS. Cell suspensions were stained with Apotracker Green dye (BioLegend) for 15 min, followed by fixation in 4% PFA for 30 mins. Cells were then pelleted, PBS washed and re-suspended in FACS buffer followed by 7-AAD (BioLegend) staining for 10 mins. Dual stained cells were PBS washed and analysed by flow cytometry using a FACSymphony (BD Biosciences). The resulting data was analysed using FlowJo v10 software (BD Biosciences).

### RNA extraction and RT-qPCR

Isolation of cellular RNAs was performed using a Direct-zol RNA Miniprep Plus Kit (Zymo Research) according to the manufacturer’s protocol. RNA quantification was performed using a NanoDrop 2000 spectrophotometer (ThermoFisher Scientific). Extracted RNAs were either used directly for reverse transcription or stored at −80*^◦^*C. RNAs were reverse transcribed into complementary DNA (cDNA) using a QuantiNova Reverse Transcript Kit (Qiagen) according to the manufacturer’s instructions. RT-qPCR of random hexamer primed cDNAs was performed using 2× Rotor-Gene SYBR Green PCR Mastermix (Qiagen) and run on a Rotor-Gene Q real-time PCR thermocycler (Qiagen). YFV genome copies in total cellular RNAs were determined using validated YFV-specific primer-pairs [78]. A YFV 17D genome containing plasmid was serially diluted to generate a standard curve to enable genome quantification.

### YFV diversity analysis

270 YFV genomes from historical and contemporary virulent strains were downloaded from GenBank (NCBI). Nucleotide sequences were aligned to ensure ORF maintenance using MEGAX[79] and percentages of variable sites in individual proteins were also calculated. Pairwise distance matrices for nucleotide and encoded amino acids were calculated in CLC genomics workbench (Qiagen). Phylogenetic reconstruction was performed using the ML method implemented in IQtree2 with the best model finder option. Statistical support for defined YFV lineages and deep tree divergences were derived using the bootstrap approach using 1,000 repetitions.

### E heterodimer structural predictions

E structures were generated using AlphaFold3 [80]. A total of five 5 models were generated with templating turned on, ranked by prediction confidence, and the model with the highest confidence was chosen for downstream visualizations. Highlighting E protein domains and mapping specific residue locations onto E predictions were performed using ChimeraX [81].

### RNA-seq

Total RNAs were extracted from PHH as described above. Quality and quantity of extracted RNAs and subsequent sequencing libraries were determined using a Fragment Bioanalyzer (Agilent). PHH sequencing library preparation was performed according to NNSR priming method [82] as described previously [83] with the following modifications. One µg of total RNA was used for mRNA isolation using the NEBNext Poly(A) mRNA Magnetic Isolation Module (NEB). An RNA fragmentation step of 5 minutes at 95°C in the buffer of the Maxima H- reverse transcriptase (RT) (Thermo Scientific) was performed before the RT reaction, which was done in the presence of Actinomycin D. After the RT, Thermolabile Exonuclease I (NEB) was used to digest the RT primer. Following RNase H (NEB) treatment the 2nd strand cDNA synthesis was performed with Klenow exo(-) (NEB). After a 1.5-volume magnetic bead purification, 10% of the sample volume was subjected to RT-qPCR to determine the number of amplification cycles required for the barcoding PCRs.

Libraries were sequenced using the NextSeq 2000 platform (Illumina) with a single-ended setting. Downstream analyses of host sequences were performed using CLC Genomics Workbench (Qiagen). Briefly, FastQ files were aligned against the Hg38 human reference genome, with annotated gene and transcript tracks. Identification of differentially expressed genes DEGs between conditions was performed with false discovery rate (FDR) correction for multiple comparisons. Gene Ontology (GO) enrichment analysis was performed with a minimum 1.5-fold change in expression threshold setting, with an FDR p-value <0.05 considered significant. Enriched GO biological process categories identified for 17D-204, AS27 or UG48 infections of PHH at each time point (24, 48 and 72hpi) were clustered based on their semantic similarity using the binary cut algorithm implemented in the simplifyEnrichment package [84]. This implementation also removed obsolete or redundant GO terms prior to clustering. Enrichment significance values were visualized as heatmaps. Volcano plots were generated using the R package EnhancedVolcano v1.22.0[85]. Volcano and GO dotplots were generated or improved with the R package ‘ggplot2’ v3.5.1.

To generate YFV consensus sequences from laboratory YFV strains, raw sequencing reads were first checked for quality using FastQC version v0.12.1 [86] followed by quality trimming with Trimmomatic version v0.40[87]. Trimmed reads were mapped to viral reference genomes (GenBank accession IDs: JX949181 [17D]; AY640589 [Asibi 1927]; and AY968065 [Uganda 1948]) using BWA-MEM version v0.7.17-r1188 [88]. The resulting contigs were sorted and indexed, followed by variant calling and consensus sequence generation using Samtools and BCFtools version v1.19[89].

Visium spatial transcriptomics and Single cell RNA sequencing data were analyzed using Seurat (v5.3.1). Raw 10x Genomics data were downloaded from the Liver Cell Atlas (https://www.livercellatlas.org), loaded into a Seurat object, and combined with existing cell type annotations, followed by data normalization and scaling. Major liver cell types were selected, and gene expression was visualized using violin plots with a uniform y-axis scale for comparison

## Data availability

RNA-seq data associated with YFV infected PHH from this study were submitted to the NCBI GEO database and can be accessed under the accession number GSE306280.

## Supporting information

ManuscriptYFV_200126Linked.pdf

## Acknowledgements

E.H. was supported by DFG grant 416701689 and the LOEWE Center DRUID (Novel Drug Targets against Poverty-related and Neglected Tropical Diseases). E.S. was supported by funding from the Horizon Europe program under grant agreement No. 101191666 for the project ‘Identification of Novel Viral Entry Factors and Development of Antiviral Approaches’ (DEFENDER). D.T. was supported by grants from the German Federal Ministry of Research, Technology and Space (BMFTR, project VirBio: 01KI2106). R.J.P.B. was supported by grants from the German Federal Health Ministry (BMG, project: ZMVI1-2516FSB416), Zoonoses Platform/BMFTR (project VIRASCREEN: 01KI2113), and intramural funding from Esther Werner (Paul-Ehrlich-Institute). We thank Ralf Bartenschlager (University of Heidelberg, Germany) for the Huh7 cells, Charles M. Rice (Rockefeller University, NY, USA) for the YFV-17Da and 17D-vGFP constructs, and Beate M. Kümmerer (University of Bonn, Germany) for the pYFV-Asibi ic construct. We also thank Michael Mühlebach, Renate König and Esther Werner (Paul-Ehrlich-Institute, Germany) for general support.

