## Supplementary material for "Yellow fever vaccine propagation in primary human hepatocytes triggers antiviral and cytolytic responses": ManuscriptYFV_200126Linked.pdf

### Supplementary Figures S1-S6

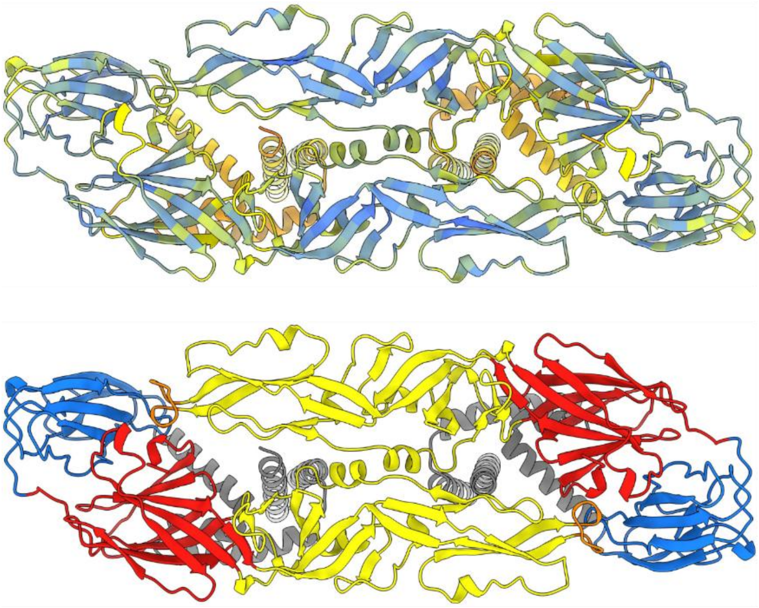

**Supplementary Fig. S1.** Confidence prediction and defined domains of YFV E proteins generated by Alphafold3. YFV E proteins are presented head-to-toe heterodimers in the prefusion conformation. Top panel. Individual residues in predicted E heterodimers are coloured according to the predicted local distance difference test (pLDDT). Dark blue: very high confidence; light blue: high confidence; yellow: intermediate confidence; orange: low confidence. Bottom panel. Predicted YFV E heterodimers with domains E-DI, E-DII, and E-DIII highlighted in red, yellow and blue, respectively. The fusion peptide is highlighted in orange, while membrane proximal and transmembrane helices are coloured grey.

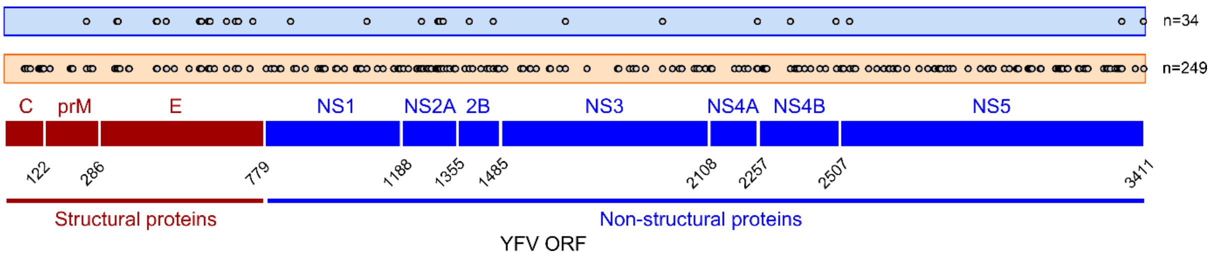


**Supplementary Fig. S2.** Location of non-synonymous mutations in genetically divergent, virulent YFV strains when compared to the 17D-204 vaccine. Plots represent comparisons of consensus sequences from Illumina sequencing of viral stocks used in this study, which were subsequently translated into amino acids and aligned. Top panel. Comparison of AS27 and 17D-204 genomes highlights a minimal set of n=34 non-synonymous mutations which differ between the virulent WAL II parental strain and the vaccine (grey circles). Middle panel. Comparison of UG48 and 17D-204 genomes identifies n=249 non-synonymous mutations which differ between the highly divergent virulent EAL strain and the vaccine (grey circles). Bottom panel. Cartoon represents the relative locations of individual YFV ORF-encoded proteins, for positional reference purposes, with encoded proteins labelled above and cleavage site co-ordinates highlighted below.

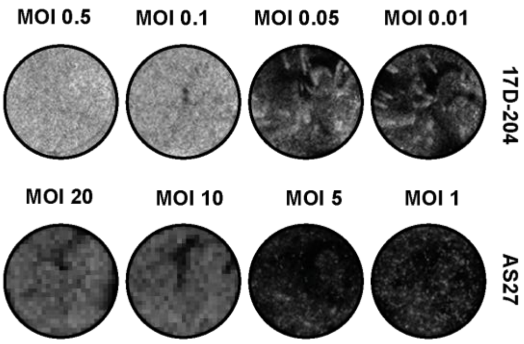


**Supplementary Fig. S3.** The 17D-204 strain shows enhanced spreading in Huh7 cells at 96 hpi, when compared to the parental Asibi strain. Huh7 cells were infected with the following virus dilutions. Top wells. 17D-204 MOI: 0.01, 0.05, 0.1 and 0.5. Lower wells. Asibi MOI: 1, 5, 10 and 20. Huh7 cells were PFA fixed at 96 hpi and stained with crystal violet.

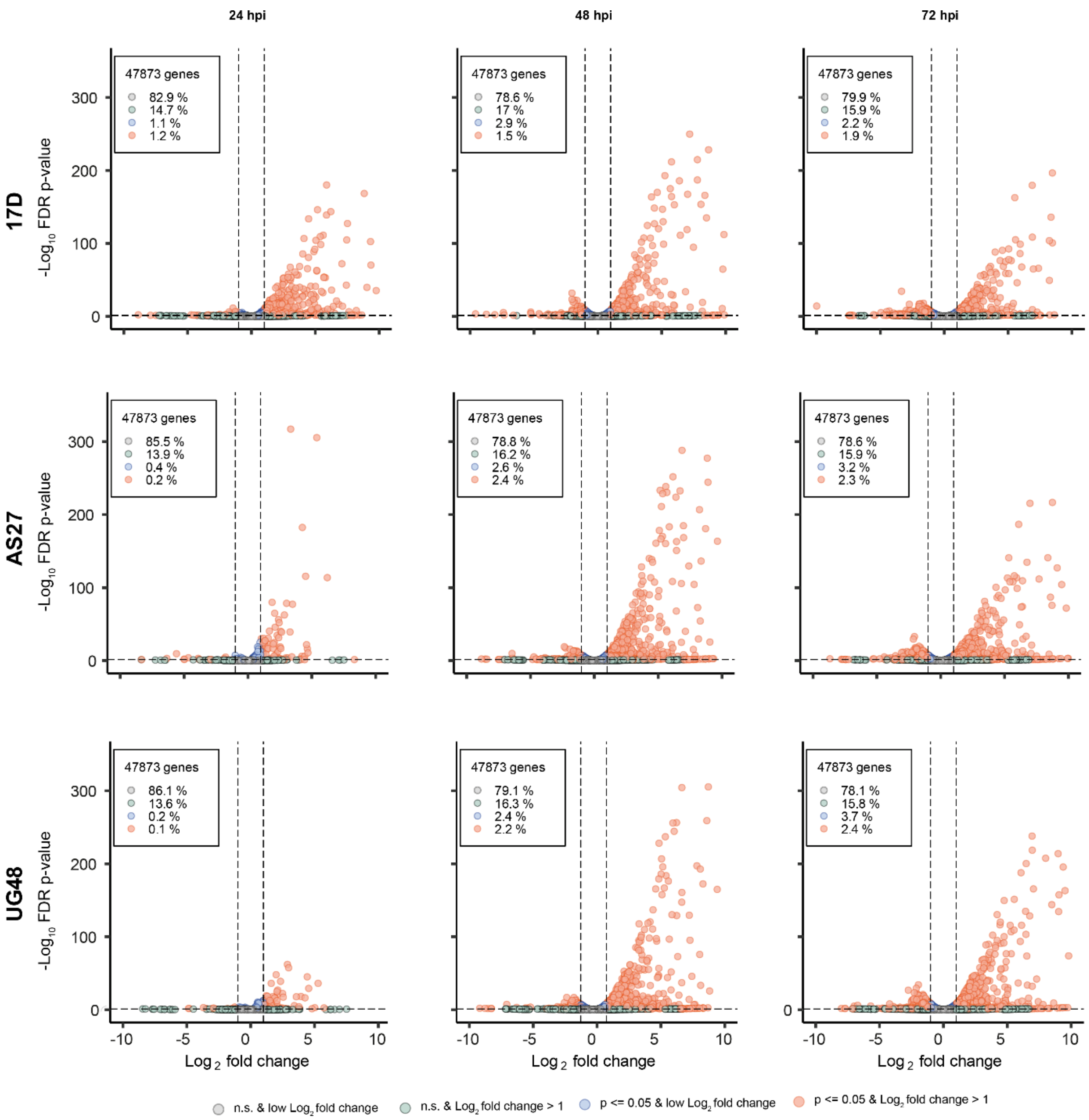


**Supplementary Fig. S4.** Volcano plots visualize DEGs induced upon infection of PHH with vaccine and virulent YFV, compared to uninfected controls. For each plot, data are derived from N=3 donors and compare PHH infected with the indicated YFV strain, to timepoint matched uninfected PHH from the same donors. Plots visualize PHH responses to infection with viruses 17D-204 (top row), AS27 (second row) and UG48 (third row) at timepoints 24 hpi (left column), 48 hpi (middle column) and 72hpi (right column). Thresholds presented in plots for FDR $\log_{10}$ *p*-values (y-axes) and $\log_2$-fold change in expression (x-axes) were 0.05 and 2, respectively. Data points represent individual genes, with genes exceeding thresholds highlighted in orange.

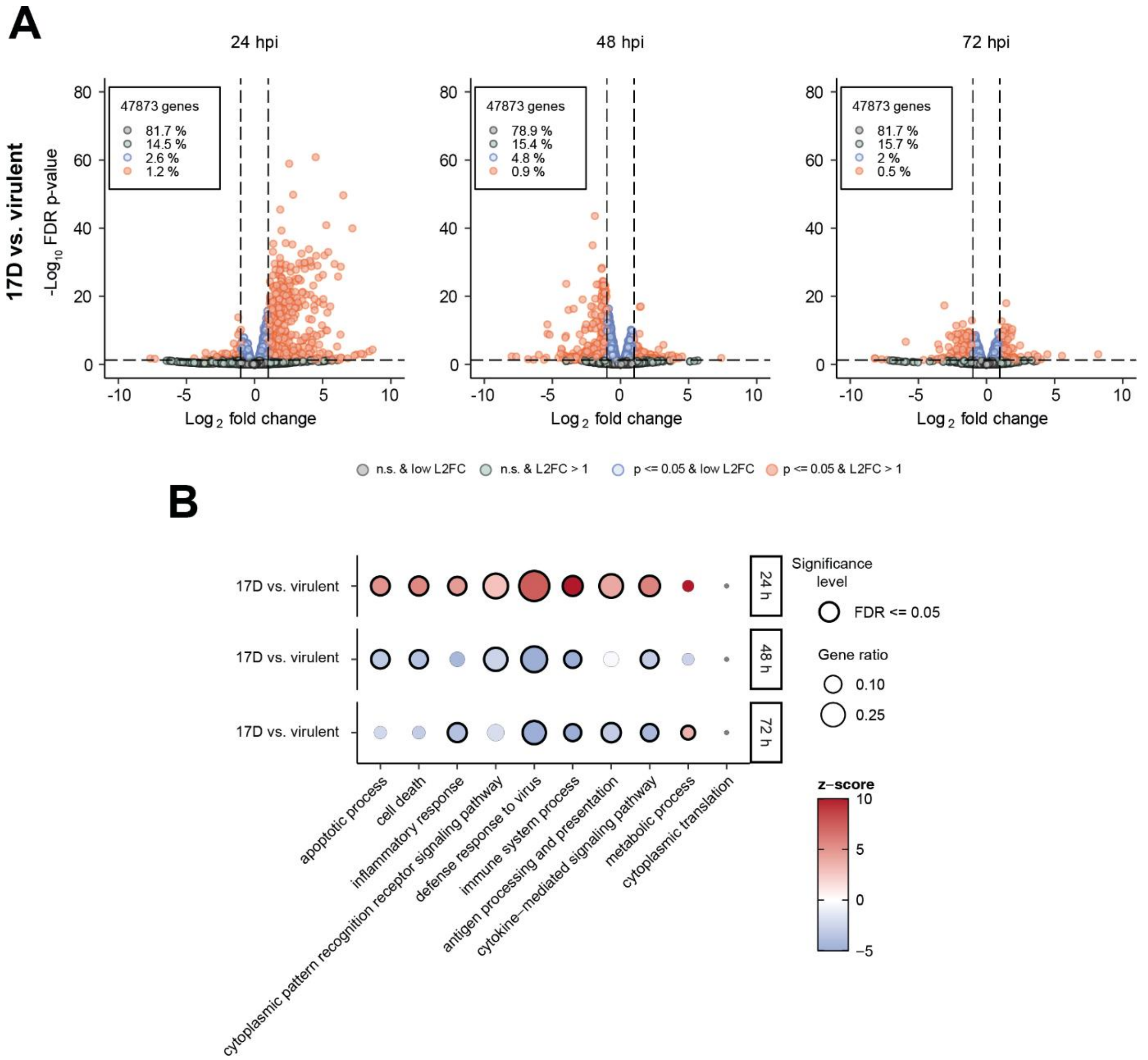


**Supplementary Fig. S5.** Plots highlight DEGs and gene programs induced by vaccine infected PHH, compared directly to virulent YFV. For each plot, data are derived from N=3 donors and compare PHH infected with 17D-204, to timepoint matched PHH infected with virulent YFV strains from the same donors. For these analyses, we tested differential expression in PHH due to 17D-204 infection, with virulent YFV-infected PHH representing the baseline control group. (A) Volcano plots visualize DEGs at 24, 48 and 72 hpi. Peak gene induction occurs earlier for 17D-204, with significantly greater gene upregulation compared to virulent YFV strains at 24 hpi resulting in enrichment of DEGs on the right side of the plot (left panel). In contrast, peak gene induction by virulent strains occurs at 48 hpi, which represent the baseline controls, resulting in a shift in dysregulation to the left side of the plot (middle panel). Thresholds presented in plots for FDR $\log_{10}$ *p*-values (y-axes) and $\log_2$-fold change in expression (x-axes) were 0.05 and 2, respectively. Data points represent individual genes, with genes exceeding thresholds highlighted in orange. (B) Enriched GO terms in YFV-infected PHH. Dot-plot visualizes significantly enriched GO categories which are shared between strains across sampling points but exhibit temporally distinct induction kinetics (10 representative categories shown). Categories are significantly enriched in 17D-204 infected PHH compared to virulent YFV infected PHH at 24 hpi. In contrast, these same categories are significantly enriched virulent YFV strain-infected PHH compared to 17D-204 at 48 and 72 hpi.

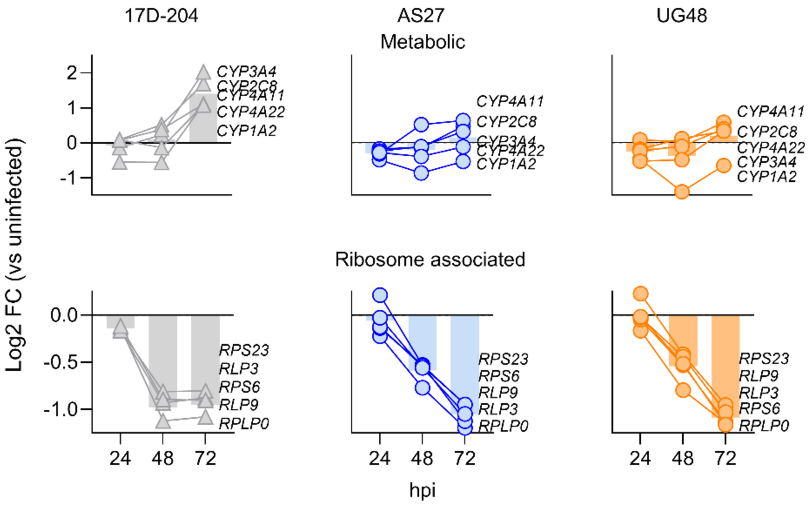


**Supplementary Fig. S6.** YFV strain-specific gene induction profiles in PHH. Plots compare induction profiles for selected categories of genes encoding gene involved in hepatic metabolism (top panels) and genes encoding ribosome associated proteins (bottom panels). Individual data points visualize gene induction ($log_2$ fold change) compared to timepoint matched uninfected PHHs and represent the mean induction for n=3 donors infected with the indicated YFV strains.
